# Facilitating CG simulations with MAD: the MArtini Database Server

**DOI:** 10.1101/2022.08.03.502585

**Authors:** Cécile Hilpert, Louis Beranger, Paulo C.T. Souza, Petteri A. Vainikka, Vincent Nieto, Siewert J. Marrink, Luca Monticelli, Guillaume Launay

**Affiliations:** Microbiologie Moléculaire et Biochimie Structurale (MMSB), UMR 5086 CNRS University of Lyon. 7 passage du Vercors, 69367 LYON, France; Groningen Biomolecular Sciences and Biotechnology Institute and Zernike Institute for Advanced Material, University of Groningen, Groningen, the Netherlands

## Abstract

The MArtini Database (MAD - www.mad.ibcp.fr) is a web server designed for the sharing structures and topologies of molecules parameterized with the Martini coarse-grained (CG) force field. MAD can also convert atomistic structures into CG structures and prepare complex systems (including proteins, lipids etc.) for molecular dynamics (MD) simulations at the CG level. It is dedicated to the generation of input files for Martini 3, the most recent version of this popular CG force field. Specifically, the MAD server currently includes tools to submit or retrieve CG models of a wide range of molecules (lipids, carbohydrates, nanoparticles, etc.), transform atomistic protein structures into CG structures and topologies, with fine control on the process and assemble biomolecules into large systems and deliver all files necessary to start simulations in the GROMACS MD engine.

## Introduction

Coarse-grained (CG) force fields allow simulations of macromolecular systems on time and length scales beyond reach for atomistic descriptions. During the past two decades coarse-graining has become a popular solution for the study of a large variety of biological problems^1,2^ as well as in materials science.^3^ This created the need for tools to facilitate the preparation and analysis of CG system. These tools are often provided as web services with graphical interfaces that require no installation by the user, while guiding him/her in the choice of the parameters and the validation of the results. Currently available web services are meeting various needs such as: supporting the generation of multiple or specific CG force field parameters, preparing the necessary files for molecular dynamics (MD) simulations, performing quick equilibration of systems or even running MD simulations for simple CG models.^4–15^

The Martini force field is one of the most popular choices among the coarse-grained force fields available,^16,17^ and offers the possibility to describe molecular interactions in systems containing lipids,^16,18,19^ proteins,^20,21^ carbohydrates,^22,23^ nucleic acids,^24,25^ polymers,^26–28^ nanoparticles,^29,30^ and other molecular systems, recently reviewed in.^31,32^ With the release of Martini 3, ^17^ many new systems and applications were also on reach of the model, including drug-like small-molecules, ^33,34^ ionic liquids,^35^ deep eutectic solvents^36^ and poly-electrolyte coacervates.^37^ Specific tools for the Martini community already exist, ^19,21,38–40^ but they are only partially covering the needs of the users, in particular the available web services. The CGMD/MERMAID webserver^14,15^ is designed the for simulation of membrane or soluble proteins but is currently limited to version 2 of the Martini force field. The CHARMM-GUI server^41,42^ allows for the preparation of MD input files for versions 2 and 3 of Martini, but offers limited possibility to edit the models.

The CG representation of a molecule in Martini is obtained by combining molecular fragments representing well-defined chemical moieties, modeled by particles or beads. A total of 843 particle types are currently available in Martini 3. ^17^ Such large number of different particle types can represent with high specificity the polarity, size, and hydrogen bonding capabilities of the building blocks they represent. Determining the most appropriate set of beads and their associated bonded parameters is a critical step in the preparation of a CG molecule. This so-called “parameterization” step often requires expert knowledge of the molecule chemistry associated with a validation protocol to ensure that thermodynamic properties can still be reproduced under the CG model. Thus the creation of accurate CG representation of molecules, while more than ever accessible, remains a daunting crafting task for most users. Although many automatic parameterization protocols have been recently developed,^43–46^ there is still a need of an extended and curated database of models that can be used as reference to access their accuracy or even be used for the calibration of such approaches. Furthermore, as the set of Martini models is growing and diversifying, the need of CG molecule repositories is becoming critical resource for the long term availability of molecular models and to make simulations reproducible.

In this context, the MAD server aims at making the set up a CG simulation with Martini accessible to the wide Martini users community, by providing the resources to obtain the CG molecules and preparing the entire system for MD simulation. To achieve this goal, the MAD server is extending the capabilities of the original Martini molecule repository (http://cgmartini.nl/) in three directions. The `MAD:Database` is storing a large collection of CG molecules readily available for MD simulations. The database is organized in a userfriendly way with modern content browsing and viewing capabilities. For macromolecules such as proteins, not present in the database, the `MAD:Molecule Builder` tool can produce coarse grained models based on its uploaded all-atom coordinates. Finally, the `MAD:System Builder` assemble many CG molecules in a simulation box and delivers all the files necessary to start the MD simulation.

Because both `MAD:Molecule Builder` and `MAD:System Builder` services make heavy usage of the server computational resources, a login is required to use them. Logged-in user are also granted access to a private storage and a job history with resuming capabilities.

## Material and Methods

### Overall Organization

The MAD ecosystem can be depicted as a database of CG molecules and CG tools connected together with computational resources and private storage for the user (see Fig.1). The main point of access to the MAD web server is its welcome page (see Fig.2), where a left menu hosts links to directly explore the database, access the tools (Molecule Builder and System Builder) or download the supported force-field files. The lower-half of the menu is for users to manage their account: profile, molecule contributed to the database and submitted jobs.

**Figure 1:**
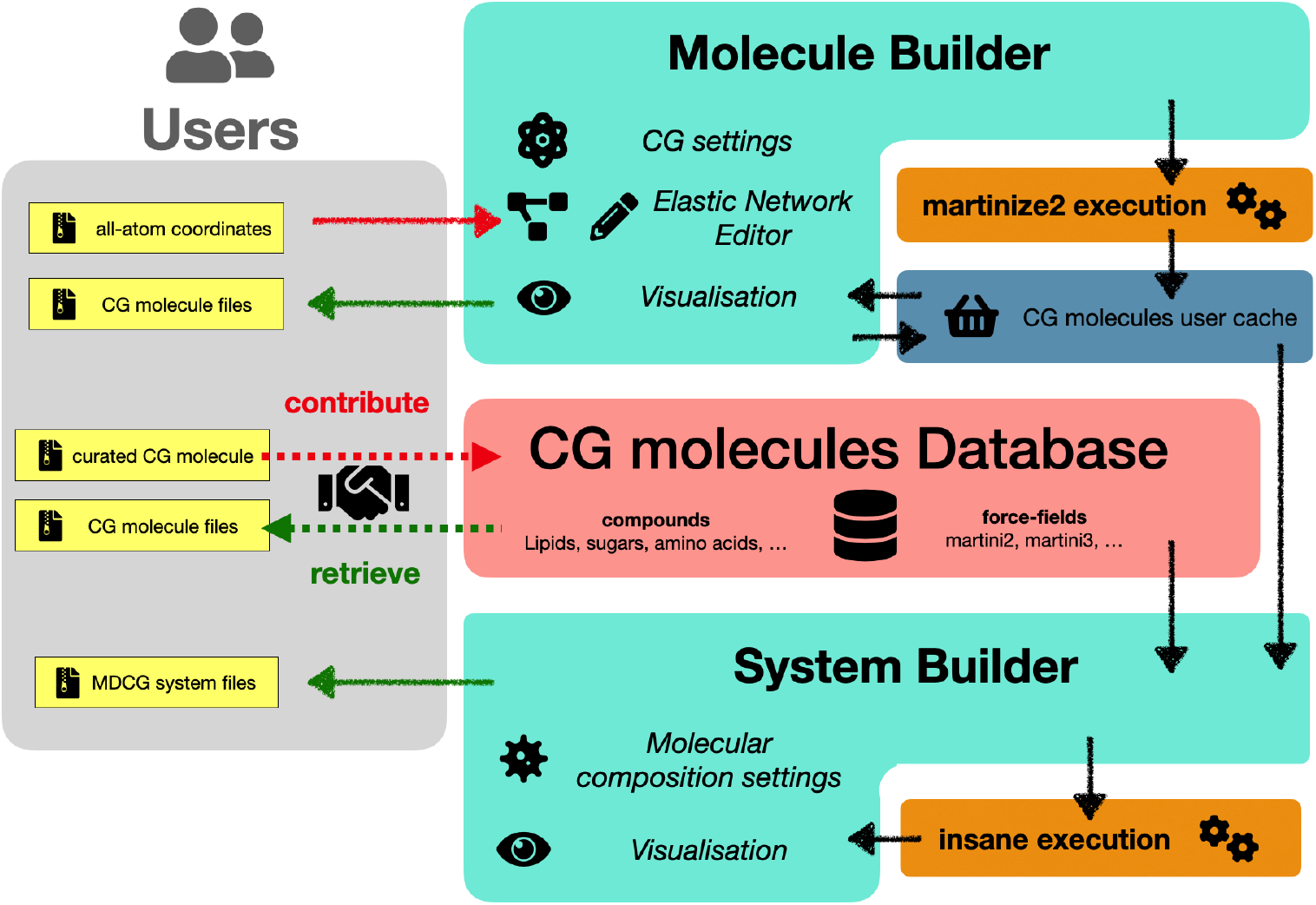
Workflow of the web server. CG models of molecules can be retrieved from or contributed to the `MAD:Database`, which covers a range of molecule types and force field versions. Alternatively, all-atom structures can be submitted to the `Molecule Builder` with control over the CG process by the *martinize2* program. Every user is granted a private storage which holds a copy of all user generated models. The `System Builder` interface can use files from the main database and from the user private storage. Submitted structures are processed by *insane* to obtain the full set of requires files to run MD.

**Figure 2:**
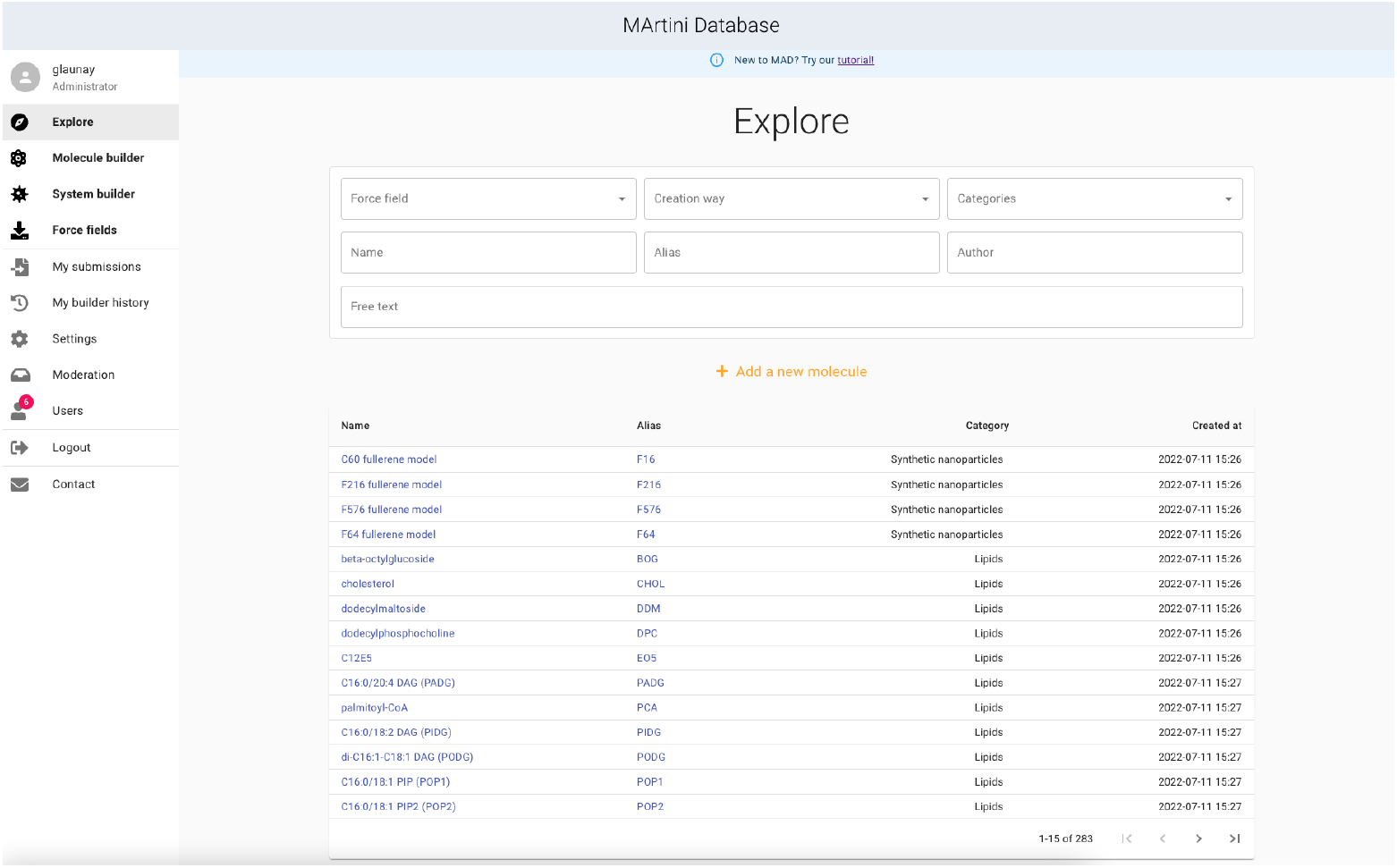
The MAD web server startup page is made of two sections. The left section is a menu dedicated to resources access and user controls. The right side is a table of the currently available CG molecules.

In fact, every registered MAD user is granted a private history module, accessible from this menu under the “my builder history” icon. Here, all his/her previous CG operations performed on the MAD server are reported to the user. Each job is labeled by its input, date of creation, the version of the targeted force field and the type of the operation. From the history panel, the user can visualize a model and eventually resume its modifications. Download links of the corresponding CG files are also available. Models can be deleted from this panel; in any case, their data will not be conserved over 15 days on the MAD file system.

The MAD server is built on a front-end to back-end architecture. The front-end, which is based on the React web component framework, carries most of the steps for the submission, validation, visualisation, and edition of structures (depicted in cyan in Fig.1). The back-end, which is based on a NodeJS/Express platform, interacts with a slurm scheduler to launch GROMACS,^47^ *martinize2* and *insane* ^19^ jobs on the MAD cluster. The public database of molecule is operated by the NoSQL Couch SGDB. All the molecular visualizations are performed by the NGL^48^ JavaScript library.

### Database

The database is designed to contain a wide variety of CG molecules which can be expanded through uploaded contributions by users. Each entry of the database corresponds to a particular association of molecule and force field version. The information of an entry is stored as a specific collection of CG files: topology files (.top and itp extension) and coordinate files (gro extension). Currently, GROMACS^47^ is the main supported MD engine, but with possible future implementation of Martini in codes as OpenMM^49^ and NAMD,^50^ more file formats may be supported by MAD. Force-field conversion tools ^51–53^ between different MD engines may be consider by the users, but features of Martini models as virtual sites may not be simply adapted in other codes.

The welcome page of the database (mad.ibcp.fr/explore) displays a top section dedicated to the custom search and a bottom section which gives direct access to the currently stored molecule in table form. Each table row links to a single molecule description page. Alternatively, molecules can be searched in the database with the top section formula of the welcome page. Here, molecules can be searched by force field name (different Martini versions are available), creation methods (manually or by different tools), biochemical category (e.g. carbohydrates, lipids, etc.) and free text searches within name, alias, or whole entry.

The search and the table browsing methods will both link to the MAD description page (Figure 3) of a molecule. The MAD molecule pages are made of four sections. The first one is the `General information` section which displays the alias, name and category of the molecule along with all the comments section extracted from the corresponding itp file. The comments can provide useful information such as the command line arguments used to generate the CG files. The top-right section of a MAD molecule page is an interactive CG molecular view of the compound, where each sphere is a Martini bead with color and size representing the bead types (Figure 3B). The `Details` section displays references for the molecules and a download link to bundled topology and structure files.

**Figure 3:**
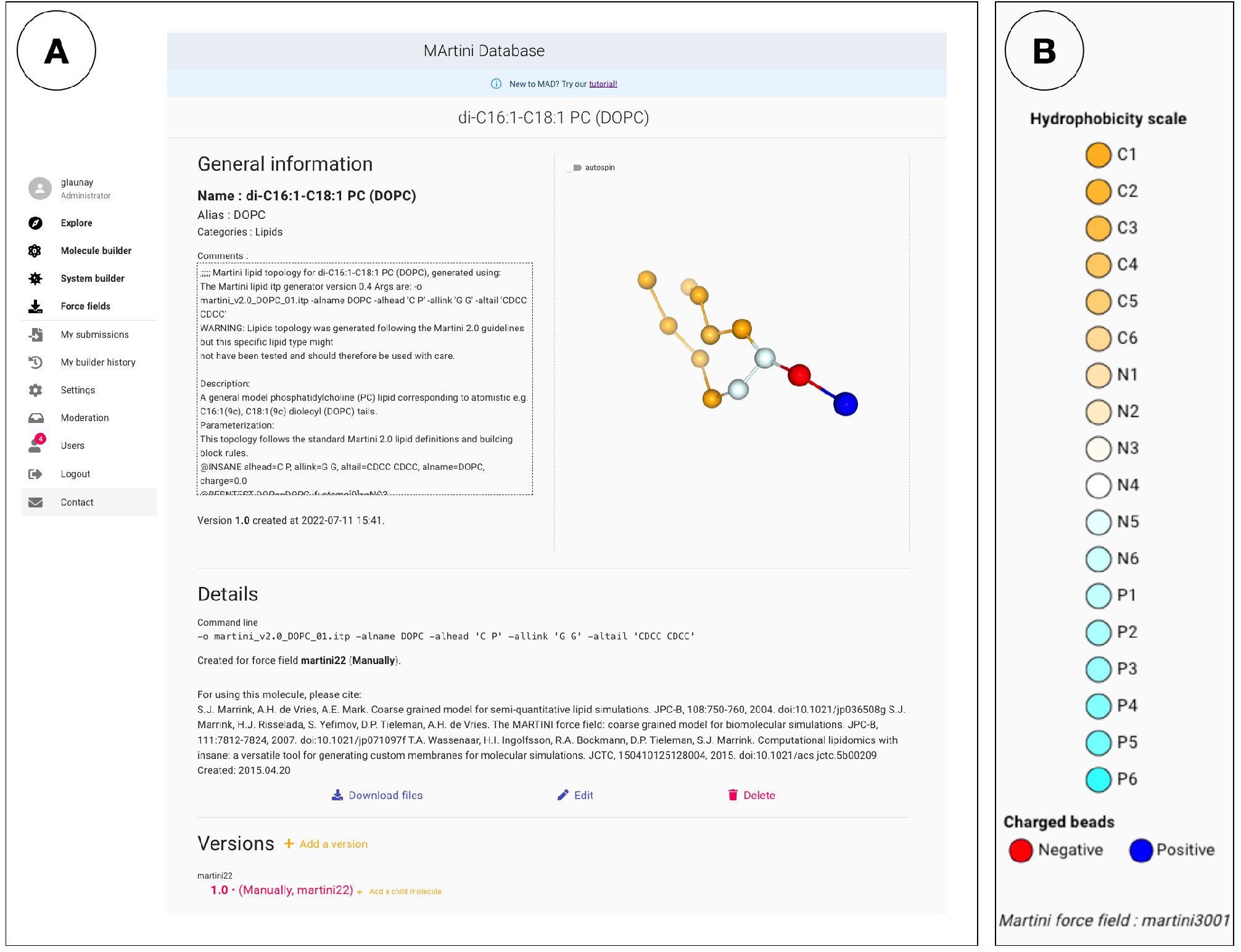
A: MAD page of dioleoylphosphatidylcholine (DOPC); B: particle color code used in MAD; the color scale broadly follows particle polarity, with red and blue colors for charged particles.

At the bottom of the page, the `Versions` section displays the trees version tracking of all the available models of the molecule. Each of these trees corresponds to a specific version of the Martini force field: only force field trees with at least one model in the database are shown. Within a tree, a node represents the specific model (and files) of a molecule stored in the database. The node of the currently displayed database entry is colored in red, the others node in the tree are links to the web page of different models of the same molecule. The children of a molecular model are models which were declared as being derived from this parent model. By clicking on the edit button, the user can effectively submit a new “child” version of the model currently displayed.

The database is open, i.e. any registered user can submit a new CG model. Submission requires the Martini files (itp, top) and at least one reference to a publication describing the derivation of the parameters. Submitted molecules will go through a quick curation process by martini developers. If the molecule is new to the database a new entry will be created. In cases where previous versions would exist, the newly submitted version will be added to the corresponding position (molecule type and force field number) in the version tracking tree.

### Molecule Builder

The `MAD:Molecule Builder` tool generates the CG structure and topology from an all-atom structure. It is currently used to coarse grain proteins, but other types of polymers will be accepted in the future. The `Molecule Builder` is built on top of our local installation of the vermouth package (https://github.com/marrink-lab/vermouth-martinize) and the *martinize2* program. It guides the user in the choice of the input parameters and provides handy edition/post-processing capabilities of the output CG structures. The interface of the `MAD:Molecule Builder` is centered around an interactive molecular representation of the user molecule. This interactive viewer is accompanied by a left-ended panel which provides the set of control commands appropriate to each molecular building stage.

As a first step, the user uploads the all-atom structure (in PDB format) to be processed. The uploaded structure is displayed in the molecular viewer, along with CG settings in the left panel. These settings control the execution of the *martinize2* program. The force fields drop-down menu features the different versions of the Martini force field available. Currently supported versions are Martini 2.2, Martini 2.2 polarized, and Martini 3.0.0.1.

The `Mode` option controls the setting of position restraints, which can be based on all-beads or backbone positions only. Bead coordinates restraints are generally useful during the equilibration simulation of the molecule, in order to maintain the protein fold. Three values of `Mode` are available: classic, elastic, ^54^ and GOMartini. ^55^ The classic mode will only use pairwise beads bonds, while elastic and Go model options respectively apply additional harmonic and Lennard-Jones potentials. The Go model contact map follows the approach of Wolek et al.,^56^ which has been implemented by Moreira et al. ^57^

The N-terminal and C-terminal fixes modify the protein terminal particles, to improve representations of functional groups charges and geometries, and are activated by default. The user may activate the “side chain fix”, which promotes protein stability and increases the reliability of the structures during MD by the addition of dihedral angles between SC1-BB-BB-SC1 beads; this provides more realistic side chain orientations.^58^ The “cystein bridges” options controls the automatic detection of cysteine residues and the application of covalent bonds between cysteine side-chains when the distance between the sulfur atoms is below a threshold of 0.216 nm. In case of long-running computations, the email toggle option makes it possible to be contacted by email upon `MAD:Molecule Builder` job completion. The email will enclose a link to access and visualize the data, which are privately stored on the server for a period of 15 days.

In case of a job failure, the user would have access to the appropriate logs. Most failures are caused by improper values in the uploaded PDB file, which can easily be traced and fixed using the provided failure logging information.

Once the structure has been coarse-grained, the user can inspect it and, most importantly, can interactively modify the set of constraints that *martinize2* created based on the chosen value of the `Mode` option. This set of constraints defines a network of elastic bond (EB), a mean to preserve the protein structure (intra-domain) while promoting molecular motions of interests (domain-domain interactions). By default, the *martinize2* software will deduce an initial network of EB from the all-atom structure. This network often needs to be edited based on (experimental) information on protein structure and dynamics. The EB editor of the `MAD:Molecular Builder` greatly facilitates this editing process. The creation/deletion of an EB requires a simple click on two beads to add or delete an elastic bond; more sophisticated modifications are achieved by the usage of a selection language to build all EB at once between two selected groups of beads. The `MAD:Molecular Builder` automatically encodes the EB network in the appropriate CG files. All modifications to the EB network can be reverted through the history section of the `MAD:Molecular Builder`.

Any stage of molecular editing in the `MAD:Molecule Builder` can be privately stashed as alternative model into the user private history; the user can access them later to resume the modifications or download the structure and topology files.

### System Builder

The `MAD:System Builder` can combine several CG structures to create large systems ready for use in MD simulations. The following types of structures are allowed: models from the `MAD:Database`, CG molecules from the user private stash, topology and structure files uploaded by the user.

The first type of system that the `MAD:System Builder` can produce are phospholipid bilayers in water solution. The builder allows to configure the lipid types and ratio, including the possibility of different compositions for the upper and lower leaflet. Salt concentration and total charge can be set by the user, and polarizable water can be used.

A macromolecule, usually a protein, in water solution constitutes the second type of system available within the `MAD:System Builder`. The macromolecule files (GRO, TOP, ITP) can be uploaded or imported from the database of compounds or from the user private stash. If no lipids are added during the setup, the resulting simulation box will comprise one instance of the provided macromolecule in a box of water molecules.

The `MAD:System Builder` can also setup CG MD files for system made of a protein embedded in a lipid bilayer surrounded by water molecules. Similarly to the previous cases, the user is guided through dedicated interface panels to define the protein, lipids and solvent of the system.

The `MAD:Molecule Builder` makes use of the *insane* software, ^19^ a powerful tool for the setup of large macro molecular systems in a simulation box. The `MAD:System Builder` assists the user with the setting of *insane* to deliver the appropriate simulation box. For all three types of systems, the `MAD:System Builder` features additional common and advanced controls with default values applicable to most cases. Advanced users may override several parameters if so needed: a useful common settings being the geometry of the simulation box. Caution should be exercised when modifications are made to the advanced settings as they could impair the simulation.

Following the computation of the system on the MAD cluster, a view of the computed simulation box is provided. Here, the left-hand panel features visualization options and a download link to the files required to start the simulation (Figure 4).

**Figure 4:**
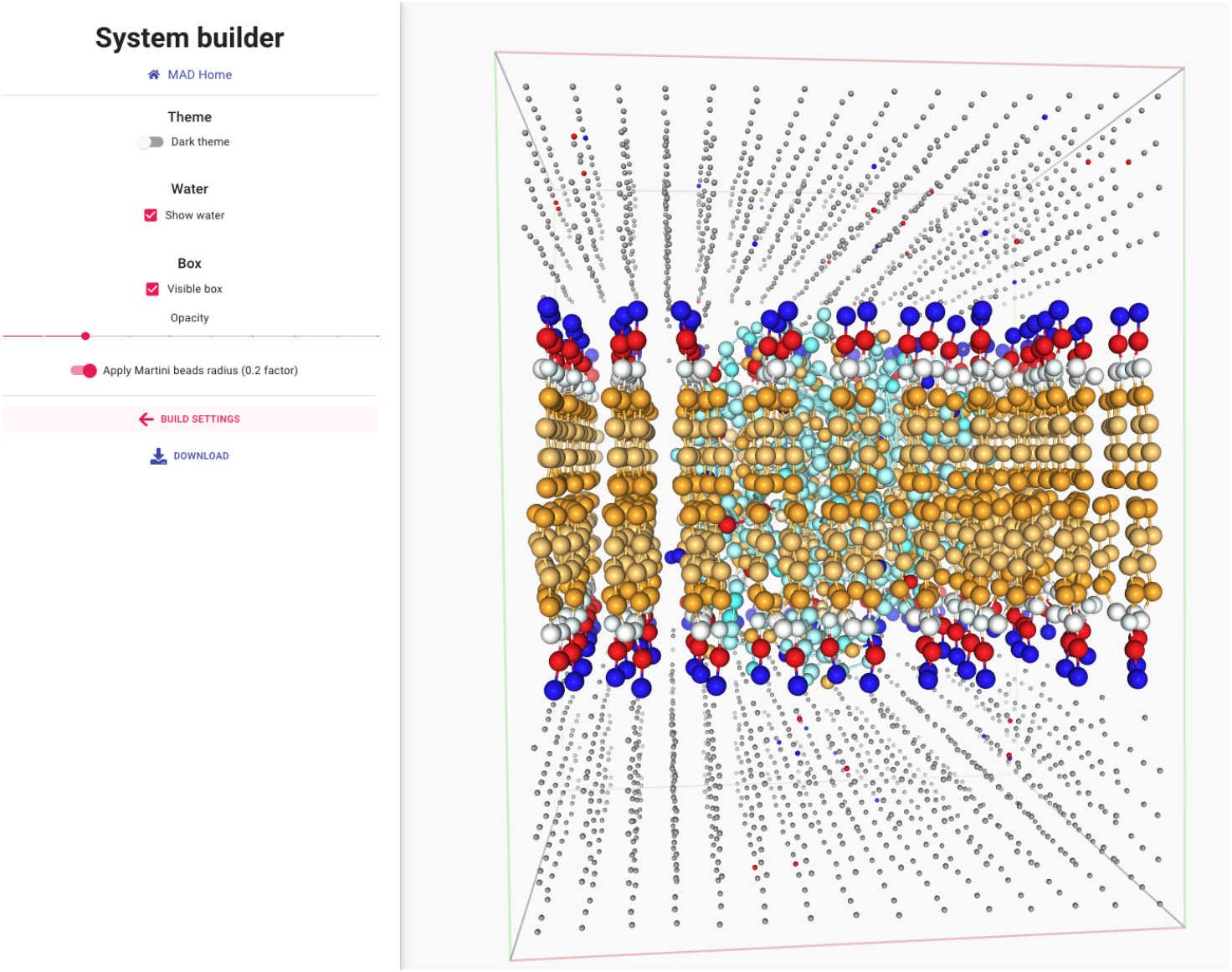
A view on the System Builder results: phospholipid bilayer and embed protein(cyan) displayed in a water-filled simulation box.

## Results and discussion

### Database content

The MAD Database currently comprises a total of 283 CG molecules belonging to the following categories: carbohydrates, polymers, amino acids, lipids, ions, phytochemical, solvents, surfactants and synthetic nanoparticles. The largest category is the lipids with currently 213 entries. Molecular entries in the database can be available for the following force fields: Martini 3.001, Martini 2.2 (for protein classic and elnedyn versions) and polarized Martini 2 versions (called 2.2P). The specific Martini 2 version developed by Monticelli and collabora-tors that is dedicated to nanoparticles and certain polymers is also included (called Martini 2.2 with CNP).^29,30,59^ Because the database is open to submission by users, these numbers are expected to change over time, with Martini 3 CG counts progressively exceeding Martini 2.

### Case study

The elastic network approach consists of a set of harmonic potential added on top of the Martini model to conserve the tertiary structure of proteins. ^54^ The network is fully dependent on the pdb structure used as reference, with the number of bonds defined by the upper and lower distance cutoff `MAD:Molecule Builder` parameters. The rigidity of the protein model is defined by the number of EB and by the force constant used. Optimal parameters for the elastic network depend of the studied protein system. It is recommended to use experimental or atomistic simulation data to calibrate the parameters of the elastic network. To illustrate the interest of the elastic network tool in `MAD:Molecule Builder`, here we show how to build models of T4 Lysozyme, a protein from the bacteriophage T4 (pdb code 181L).^60^ Once the structure file is open in the `MAD:Molecule Builder`, the force field is set to “martini3001”. With “Elastic” activated, the default setup provides for a force constant of 700 kJ/(mol.nm2), with the lower and upper cutoffs at 0.5 and 0.9 nm respectively. Finally, neutral termini, auto assignment of cysteine bridges and side-chain fix are applied (Figure 5A). Upon completion of the CG process, the resulting model is displayed (Figure 5B) using the same beads color scales as the database of compounds (Figure 3B). The automatic construction of elastic network from the all-atom structure can lead to artefact EB between CG beads. Within the `MAD:Molecule Builder`, visual inspection of the EB network of T4 Lysozyme protein superimposed onto its CG model facilitates the identification of such a case. Indeed, an abnormal EB is found to be present between T21 and Q141 (Figure 5C). As the two amino acids are distant in structure, their EB may greatly impair the flexibility of the CG model. To better prepare the model for subsequent MD simulations it recommended to delete the bond by directly clicking to select and remove (Figure 5C). Alternatively, rigidity can be added to the model in the N-terminal subdomain as exemplified in Figure 5D) where all possible EB between two amino acids selections: D10 (violet color) and D20-T21 (yellow color), are created in a single click.

**Figure 5:**
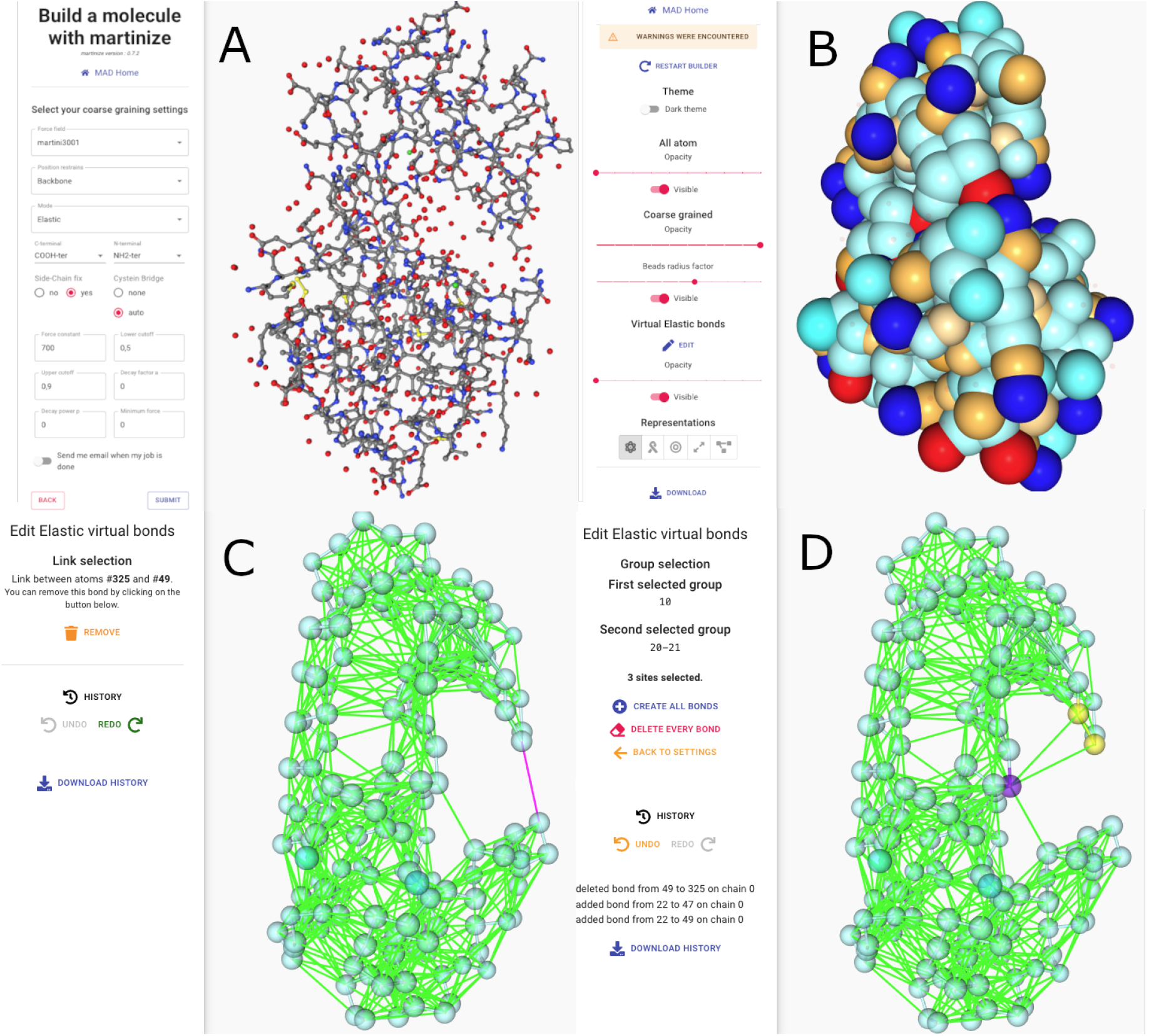
Sequential views of the process of coarse-graining the bacteriophage T4 Lysozyme (pdb code 181L): from the initial settings of parameters on the all-atom structure (A) to the visualization of the beads at a 0.6 scale on the CG structure (B), the direct (C) selection of an EB to be removed and (D) the query based selection of amino acids to connect through newly created EB. Green bonds on panels C and D do not represent covalent bonds but EB between CG beads.

### Conclusion and perspectives

We presented here the MAD server, a new web resource dedicated to the preparation of MD systems with the Martini coarse-grained force field. The MAD server provides a large collection of CG molecules ready to be used. Newly parameterized and published molecules can be uploaded to the `MAD:Database`. For molecules not yet published, topology and structure files can be provided from the user’s computer. Alternatively, all-atom structures can be coarse-grained by the `MAD:Molecule Builder`. In a final step, CG molecules from any of these sources can be combined by the `MAD:System Builder` to produce topology and structure files MD-ready. We strongly encourage users to contribute to the repository by uploading their favorite models. For the new users, a tutorial is available at https://mad.ibcp.fr/tutorial/. We plan on expanding the capabilities of the MAD server through the future integration of algorithms useful to the MARTINI users community, as Polyply.^40^

## Acknowledgement

We thank all external users for testing MAD during its development. Luca Monticelli thanks the French National Institute of Health and Medical Research (INSERM) for the support. Paulo C. T. Souza acknowledges the supported by French National Center for Scientific Research (CNRS). Further funding of Paulo C. T. Souza and Luca Monticelli came from a research collaboration with PharmCADD. Siewert J. Marrink acknowledges funding from the European Research Council via an Advanced grant Advanced Grant (COMP-O-CELL).

## Notes

https://www.mad.ibcp.fr

### Competing Interest Statement

The authors have declared no competing interest.

### Summary of Updates

fixed some typos and corrected submitted title

